# Microbial turnover and dispersal events occur in sync with plant phenology in the perennial evergreen tree crop, *Citrus sinensis*

**DOI:** 10.1101/2022.02.10.480024

**Authors:** Nichole A. Ginnan, N. Itzel De Anda, Flavia Campos Freitas Vieira, Philippe Rolshausen, M. Caroline Roper

## Abstract

Emerging research indicates that plant-associated microbes can alter plant developmental timing. However, it is unclear if host phenology impacts microbial community assembly. Microbiome studies in annuals or deciduous perennial plants face challenges in separating effects of tissue age from phenological driven effects on the microbiome. In contrast, evergreen perennial trees, like *Citrus sinensis*, retain leaves for years allowing for uniform sampling of similarly aged leaves from the same developmental cohort. This aids in separating phenological effects on the microbiome from impacts due to annual leaf maturation/senescence. Here we used this system to test the hypothesis that host phenology acts as a driver of microbiome composition. *Citrus sinensis* leaves and roots were sampled during seven phenological stages. Using amplicon-based sequencing, followed by diversity, phylogenetic, differential abundance, and network analyses we examined changes in bacterial and fungal communities. Host phenological stage is the main determinant of microbiome composition, particularly within the foliar bacteriome. Microbial enrichment/depletion patterns suggest that microbial turnover and dispersal were driving these shifts. Moreover, a subset of community shifts were phylogenetically conserved across bacterial clades suggesting that inherited traits contribute to microbe-microbe and/or plant-microbe interactions during specific phenophases. Plant phenology influences microbial community composition. These findings enhance understanding of microbiome assembly and identify microbes that potentially influence plant development and reproduction.

## INTRODUCTION

Plant phenology, the periodic timing of plant life cycle events, is innately linked to exogenous climatic variables that affect plant development, such as temperature, photoperiod, nutrient and water availability, as well as other abiotic and biotic factors^1,2^. Additionally, endogenous genome-encoded factors such as dynamic internal photosynthate source-sink pathways, intricate phytohormone signaling networks and other developmental regulatory processes mediate the transition between phenological stages^3,4^. The timing of specific developmental stages, such as flowering, can determine a plant’s geographic distribution range as well as determine crop yield and productivity^5,6^. Alterations in plant phenology can also have a cascading effect on the fitness of organisms that depend on those specific plants for nutrient acquisition, such as pollinator species^7–9^.

Citrus is a significant economic crop and provides several health benefits because of the myriad of nutrients, antioxidants, vitamins, minerals and dietary fiber found in fresh and juiced citrus fruits^10–12^. Citrus varieties are grown across the globe, and because of this, citrus phenology is well characterized to guide management strategies of different varieties for specific climatic conditions. Phenological modeling of citrus has focused primarily on buds, flowers and fruit and is used to predict bloom time across different growing regions^13^. This has implications for protecting flowers from floral pests and pathogens by allowing growers to time spray applications in an informed manner. Citrus flowers are a significant source of nectar related to honey production, particularly in California’s Central Valley. As such, bloom timing models are also important for the beekeeping industry^14^. In addition, bloom time and duration models can be extrapolated to predict fruit set^15^ and these performance models can provide yield predictions.

Soil and rhizosphere microbiomes can drive changes in flowering time in the herbaceous perennial plant *Boecheria stricta*, a wild relative of Arabidopsis^16,17^ and affect other above ground plant traits in the annual plant system, *Brassica rapa*^18,19^. However, questions about microbiome shifts associated with transitions between phenological stages have not been addressed in perennial trees, particularly domesticated evergreens like citrus^20^. Citrus phenology models primarily take into account temperature and number of degree days above a certain threshold temperature^15^, but to the best of our knowledge, have not incorporated studies on the microbial communities associated with transitions across phenological stages. The citrus microbiome is an emerging prototype for understanding microbial contributions to plant health in a perennial arboreal crop system^21–25^. Due to its well-defined phenology, citrus is an ideal system to investigate the interplay between host phenology and microbial community composition.

Several seminal studies in annual and short-lived perennial plants have characterized changes in rhizosphere and root microbiome composition across plant developmental cycles, suggesting that host phenology drives these alterations^26–31^. However, Dombrowski *et al*. 2017 suggests that initially microbiota are sequentially acquired resulting in community changes as the host ages, but eventually the microbiome matures and stabilizes, functioning independently from host development^32^. Another recent study supports the idea that time is a stronger predictor of microbiome composition than plant developmental stage^33^. This prompts discussion on whether these community shifts are a consequence of tissue age and a microbiome maturation process, or if these changes are driven by plant phenology. In addition to producing and maturing leaves and roots throughout the year, long-lived evergreen perennial plant systems retain mature leaves for up to four years, which allows for selection of leaf tissues of similar age and developmental cohort across phenophases. Because of these features, we utilized this system to help decouple tissue age from host phenological effects and tested the hypothesis that host phenology acts as a driver of community compositional shifts within the above (foliar) and below (root) ground microbiomes of citrus. Indeed, we determined that the significant shifts in both diversity and composition of the microbial community structure were primarily driven by host phenological stages and not exogenous environmental factors such as rainfall, hours of irrigation or temperature. Foliar communities were more affected by host phenology than root microbiomes, which were comparably more stable. Interestingly, major alterations in foliar microbial community composition correlated with the shifts in source-to-sink pathways of carbohydrate transit, namely during the transition from floral bud break to full flowering to fruit set. More specifically, subsets of these taxa displayed temporal turnover patterns indicating that specific taxa were enriched as trees shifted to reproductive growth associated with fruit production. We also observed taxa typically associated with pollinator species that were substantially enriched only during flowering, suggesting that these microbes were introduced into the foliar microbiome as microbial immigrants via an insect-mediated dispersal mechanism.

In agricultural plant systems, comprehensive microbiome studies allow researchers to place an emphasis on how the microbiome as a whole function to promote overall plant health by a variety of mechanisms, such as enhancing nutrient uptake or resisting pathogen ingress to promote a sustainable agroecosystem. Uncovering links between plant phenology and shifts in microbiome structure is the first step towards a mechanistic understanding of microbiome resilience over cyclical development in a perennial plant host. In addition, this can further serve as the foundation to understanding how the microbiome responds to changes in host development and, in turn, if microbiome community structure can influence host phenological transitions.

## MATERIALS AND METHODS

### Sample collection and field sites

In this longitudinal study we collected leaf (n=159) and root (n=159) samples from eight trees, at 20 timepoints (monthly for first sample year, targeted sampling second year) starting July 2017-April 2019 from Late Navel Powell sweet orange trees grown at UC Lindcove Research and Extension Center in Exeter, CA. We sampled approximately one-year old leaves that were from the previous year’s flush and, thus, in a separate developmental cohort than the reproductive structures on the trees at the time of sampling. Trees were planted in 1997 (20-22 years old at time of sampling) and managed with conventional farming strategies similar to industry orchards. Prior to each monthly sampling, trees were visually assessed, and developmental stages were recorded. Seven major phenological stages were used for categorization in this study. These included Spring vegetative shoot flush, referred to as “flush” (F) (February – March), early floral bud break and development referred to as “floral bud development” (FB) (March), full flowering (FF) (April), fruit set (FS) (May – July), exponential fruit growth and development, referred to as “fruit development” (FD) (August – October), color break (initiation of fruit maturation, CB) (November – December), and mature fruit (MF) (January) (Fig. **1**). Mature fruits were harvested between our April FF and May FS sampling events. Fertilizer treatments and/or amendments and number of hours irrigated were collected as monthly metadata variables. Each tree was divided into 4 quadrants (north, south, east, and west) and stems with attached young, fully expanded metabolically active, mature leaves from the current year (petiole attached) were collected from each quadrant and pooled into a sterile 24 oz. stand-up whirl-pak bag (Nasco B01401, Fort Atkinson, WI). Leaves were sampled from fruit bearing and non-bearing branches at random and all leaves from a single tree were pooled. Feeder roots were sampled from two sides of the tree approximately 0.5 meters away from the base of the trunk near the irrigation line and sealed in an additional sterile 24 oz. stand-up whirl-pak bag (Nasco B01401, Fort Atkinson, WI). Gloves were changed and clippers and shovels were sterilized with 30% household bleach between samples. All samples were immediately placed on ice in a cooler for transit to the laboratory, then immediately frozen at -20°C. Samples were inspected by the California Department of Food and Agriculture according to California citrus quarantine protocols prior to overnight shipment to UC Riverside on dry ice. Samples were kept frozen on dry ice while processing for downstream DNA extractions. Leaf tissue was removed from stems and chopped into smaller pieces and root tissue was rinsed with sterile water to remove surface soil. Both tissues were put in 50 ml falcon tubes and stored at -80°C then lyophilized with a benchtop freeze dryer (Labconco FreeZone 4.5L, Kansas City, MO) for 16 to 20 hours. DNA extractions were performed according to published protocols^21^. The DNA was stored at -20°C until utilized for bacterial and fungal Illumina library construction.

**Figure 1.**
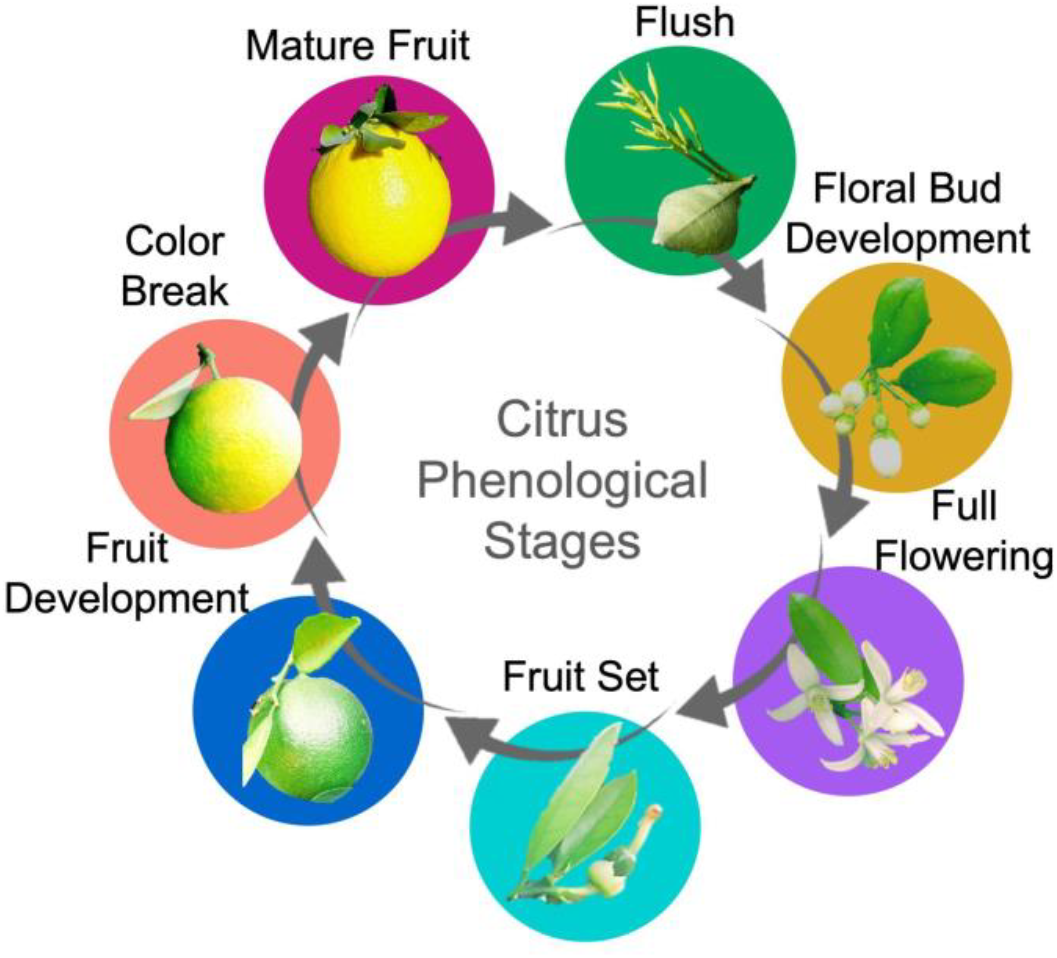
Citrus phenological stages. Cyclic seasonal development of *Citrus sinensis*.

### High-throughput sequencing library preparation

Leaf and root bacterial communities were sequenced from all leaf and root samples (20 time points). Bacterial Illumina Miseq libraries were built by amplifying the bacterial rRNA 16S V4 region using the 515FB/806RB primer set and the standardized Earth Microbiome Project protocol, which can be found online at http://press.igsb.anl.gov/earthmicrobiome/protocols-and-standards/16s/ ^34^. To limit the amplification of plant mitochondrial and plastid 16S regions, we used pPNA and mPNA clamps, which bind to these plant sequences and block binding of the bacterial 515FB/806RB. Leaf sample reactions received 0.75 μM of pPNA and mPNA clamps and 0.75 μM of mPNA was added to each root sample reaction.

The first 14 consecutive monthly leaf and root samples were included in fungal community libraries. Our preliminary analyses shifted our main focus to bacterial communities, therefore we did not sequence the fungal communities of leaf/roots collected at the last 6 time points in the second sampling year. Fungal Illumina MiSeq libraries were built by amplifying the fungal ITS1 region using the ITS1f/ITS2 primer set and the standardized Earth Microbiome Project protocol, which can be found online at http://press.igsb.anl.gov/earthmicrobiome/protocols-and-standards/its/.

Triplicate PCR reactions for each sample were pooled and amplification was verified on a 1% agarose gel. Amplicon samples were quantified for DNA concentration using Quant-iT PicoGreen (Invitrogen). Equal amounts of amplicons (240 ng) from each sample were pooled and AMPure XP (Beckman Coulter) beads were used to clean the sample library. The cleaned library was then quantified using Qubit (Qiagen) (260/280). The libraries were diluted to 20-30 μg/mL and a final quality assessment was done with a Bioanalyzer at the UCR Genomics Core facility. Paired-end sequencing (2×300) was performed on an Illumina MiSeq platform with a 20% PhiX spike included before sequencing.

### Data processing and statistics

Demultiplexed, PhiX reads removed, and Illumina adapter trimmed sequences were received from the UCR Genomics Core. Amplicon sequencing raw reads for 16S rRNA genes and fungal ITS2 region are available on the NCBI SRA database under BioProject accession number PRJNA685913. Bacterial and fungal reads were pre-processed using a USEARCH(v9.1.13)/VSEARCH(v2.3.2) pipeline. Forward and reverse sequencing files were joined with USEARCH allowing for staggered ends and up to 10 mismatches. After quality filtering using VSEARCH, there were 65.5million 16S reads and 21.1 million fungal reads. Sequences were dereplicated, singletons were removed, and OTUs were formed using USEARCH with a 97% similarity cut-off. The bacterial library produced 19,626 OTUs which were assigned to 4,183 taxonomic names using the RDP 16S database, v18. The fungal library produced 44,447 OTUs which were assigned to 31,056 taxonomic names using the UNITE fungal reference database, v02.02.2019. On average, 12.7% of the 16S library reads from each leaf and root sample were assigned as bacteria. See Supplementary Table S1 for all 16S bacterial read counts of individual samples. The remaining sequences, which were removed, were attributed to chloroplast, plant mitochondria, Archaea, or could not be assigned to a Kingdom. Our fungal libraries did not have non-specific binding issues. USEARCH was also used to create phylogenetic tree files in Newick format.

Pre-processed taxonomically assigned OTU tables were imported into R (v3.6.0). Samples with less than 1,000 reads were removed. Reads were rarified to even depth for each alpha diversity comparison. Alpha diversity was compared using the number of OTUs observed^35^. A ranked sums analysis of variance statistical test, Kruskal-Wallis, followed by a pairwise Dunn’s test with Holm’s correction for multiple comparisons were used to calculate *P*-values.

Beta diversity analyses were performed using R packages phyloseq (v1.28.0) and vegan (v2.5.6)^35,36^. Pre-processed reads were transformed using total sum scaling normalization. Using the ordinate() and plot_ordination() functions a Principal Coordinate Analysis (PCoA) was done on weighted Unifrac distances, which accounts for relative relatedness and quantitative variance of communities. Ninety-five percent confidence ellipses were added to further examine groups using the stat_ellipse() function. A permutational multivariate analysis of variance (PERMANOVA) statistical test was performed on weighted Unifrac distances using the vegan::adonis() function, including the following covariates and interaction terms in the model: adonis(dist.matrix ∼ phenology*Sample_year + Fertilizer_app + Mean_Temp*hrs_irrigated + Mean_Temp*Total_rain, permutations=999) Pairwise PERMANOVA with FDR correction was accomplished with RVAideMemoire::pairwise.perm.manova() (v0.9.74)^37^. Core microbiota identifications were performed using microbiome::core() (v1.6.0) with prevalence set at 0.75 and detection set at 0.01/100^38^. The core root bacteriome was also defined with prevalence set at 0.75 and detection set at 0.1/100, which did not impact the interpretation of the results, but greatly improved readability of the phylogenetic tree.

To understand if phylogenetic relationships impact microbial associations with phenological stages, core genera taxonomic assignments were input into a phylogenetic tree generator, phyloT v2 (https://phylot.biobyte.de/index.cgi), to generate a Newick format phylogenetic tree. Tree and metadata were visualized using the interactive tree of life visualization program, iTOL (https://itol.embl.de/)^39^. Genera with less than 50 reads were filtered out and differentially abundant populations at the genus level were identified using DESeq2 (v1.24.0) to run a parametric fit for dispersion on a negative binomial generalized linear model, followed by a Wald test with FDR adjustment to produce *P*-values^40^. Input root and leaf bacterial OTU tables used in DESeq2 analysis were rescaled using a pseudocount of 1. All possible phenological stage pairwise combinations were tested using results() and the contrast option. Using ggplot2 (v3.2.1) and phyloseq::subset_taxa() function, the relative abundance of specific species were plotted as boxplots^35,41^.

The top 300 most abundant leaf bacterial OTUs were input into a network analysis using Sparse Inverse Covariance Estimation for Ecological Association Inference, Spiec-Easi (v1.0.7), which infers interactions using neighborhood selection and the concept of conditional independence, rather than a standard correlation or covariance estimation^42^. With set.seed(1244), neighborhood modeling (mb) was executed with the nlambda set to 70 and rep.num set at 99, standard settings were used for all other parameters. Spiec-Easi results were converted to igraph format and imported into Gephi (v0.9.2). The network was visualized using a Yifan Hu layout and taxa were filtered to focus on differentially abundant taxa and immediate neighbors. Betweenness centralities were calculated using Gephi network diameter statistics and centralities were normalized to a 0-1 scale. Taxa were further filtered by abundance with the final figure showcasing the 155 most abundant OTUs within the above parameters. This significantly increased readability without impacting the interpretation of the results.

## RESULTS

### Significant shifts in alpha diversity occur across phenological stages

We focused our study on seven citrus phenophases that included, Spring vegetative shoot flush, referred to as “flush” (F), early floral bud break and development (FB), full flowering (FF), fruit set (FS), exponential fruit growth and development, referred to as “fruit development” (FD), color break (initiation of fruit maturation, CB) and mature fruit (MF). Citrus phenological stages can overlap on individual trees and some stages span multiple months, thus, some stages include multiple months of sampling (Fig. **1** and Supplementary Fig. **S1**). Overall, bacterial and fungal leaf microbiomes had the most significant shifts in alpha diversity across phenological stages when compared to the root microbiomes. Specifically, alpha diversity in both the leaf bacteriome and mycobiome remained consistent as trees transitioned from leaf flush to flowering (floral bud development and full flowering). Following full flowering, there was a significant increase in alpha diversity in the leaf bacteriome and mycobiome at fruit set (Fig **2a,b**). Species richness within the leaf bacteriome significantly decreased when trees transitioned from fruit growth and development to color break and mature fruit stages.

**Figure 2.**
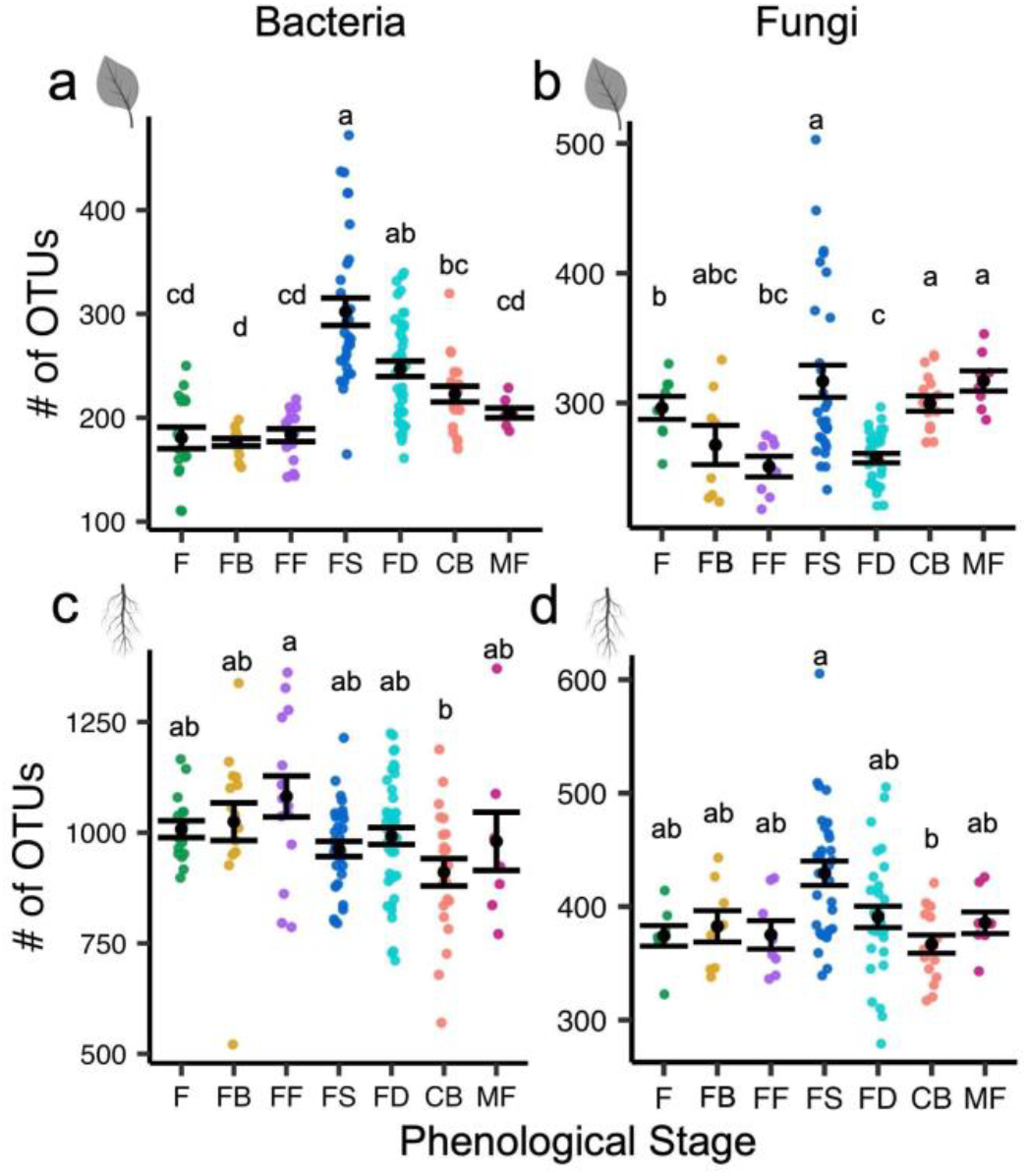
Species richness varies across phenophases. Alpha diversity plot of richness (No. of OTUs) for individual sampling events for bacterial leaf (a), fungal leaf (b), bacterial root (c), and fungal root (d) communities. Black points, “⋅”, represent the mean and error bars represent standard error. Letters indicate a significant difference of *P* ≤ 0.05, determined using a Kruskal-Wallis test, with a pairwise Dunn’s test and correcting for multiple comparisons with Holm’s method. Phenological stages on the x-axis include: flush (F), floral bud break (FB), full flowering (FF), fruit set (FS), fruit development (FD), color break (CB) and mature fruit (MF).

Despite being relatively stable across the study, root bacteriome alpha diversity peaked during full flowering. Similar to the overall leaf microbiome, the root mycobiome had the highest alpha diversity during fruit set (Fig. **2c, d**). Our study did not discriminate between rhizoplane and endophytic root microbiota nor was it possible to select feeder roots of a specific age cohort. Future work that separates these compartments in similarly aged roots may reveal more finely resolved shifts in species richness associated with these root environments.

### Host phenological stage is a major determinant of community composition

Although climatic variables can be difficult to uncouple from plant development variables, the greatest amount of the variation in the data was attributed to the host phenological stage for all four communities (PERMANOVA, *P* ≤0.001, *r*^*2*^ = 0.081 - 0.280) (Table **1**, Fig. **3**). Time (i.e., sample year) had less impact than phenology on beta diversity across all communities (Table **1**, Supplementary Fig. **S2**). Interestingly, the community composition (beta diversity) of leaf bacteriome and mycobiome was more influenced by host phenology than root communities, indicating that changes in host phenology had a larger influence on diversity within foliar microbiomes than in root microbiomes. Specifically, a principal coordinate analysis of UniFrac distances indicated significant clustering of individual microbial communities by phenological stage (Fig. **3**). In a pairwise comparison of community compositional differences between each phenological stage, the leaf bacteriome had the greatest number of significant adjusted *P*-values, with 21 out of the 21 pairwise comparisons being significantly different, followed by the leaf mycobiome (19 out of 21 comparisons) (Supplementary Table S2). Root communities had fewer significantly different pairwise comparisons (Root Mycobiome = 5/21, Root Bacteriome = 8/21).

**Figure 3.**
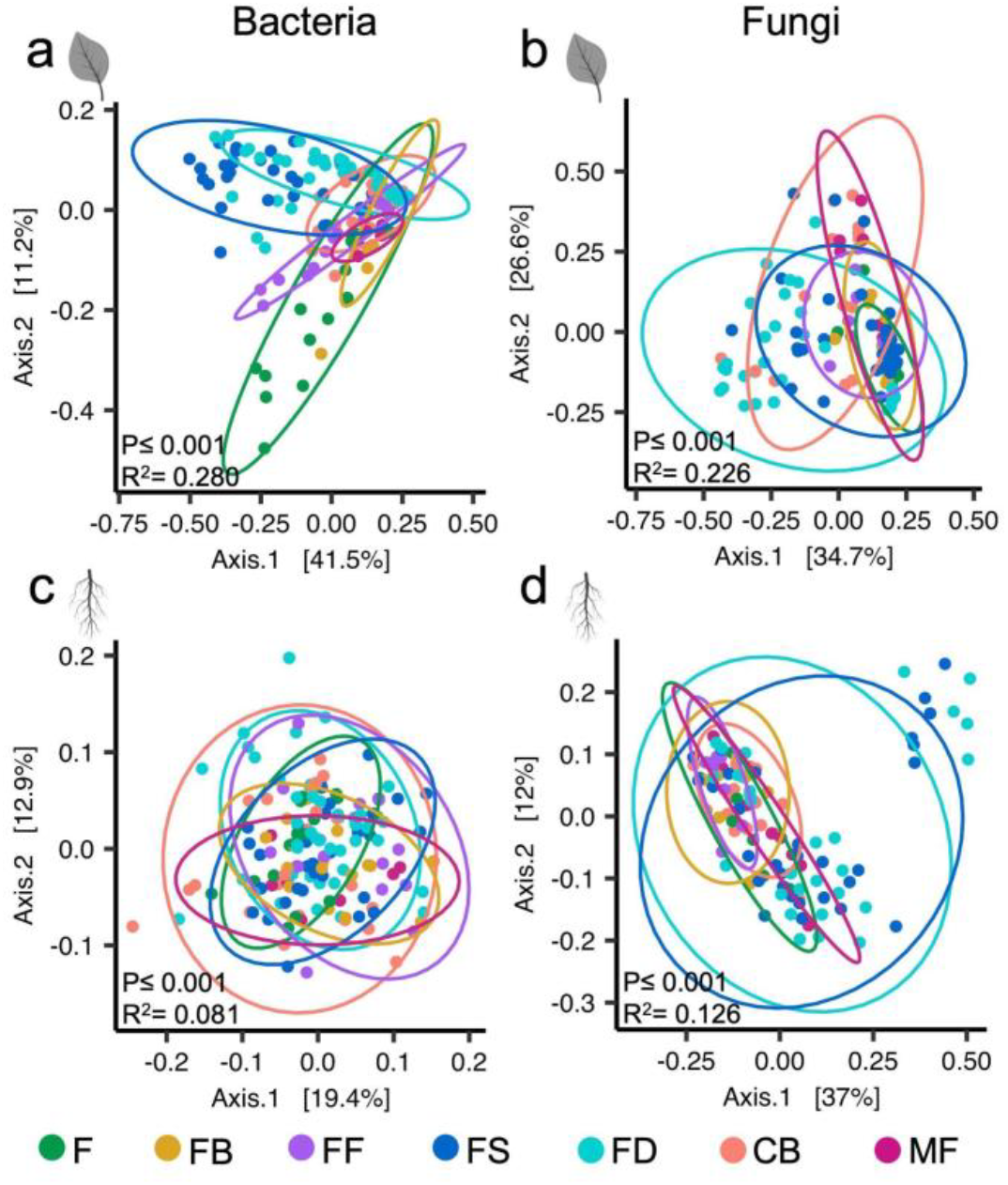
Host phenology affects community diversity and composition. Beta diversity PCoA plots of bacterial leaf (a), fungal leaf (b), bacterial root (c), and fungal root (d) communities. Points are colored by phenological stage and represent a complete community from a single leaf or root sample. Ellipses represent 95% confidence intervals. The *P*-values and *r*^*2*^ values were obtained using a PERMANOVA (Adonis). Phenological stages include: flush (F), floral bud break (FB), full flowering (FF), fruit set (FS), fruit development (FD), color break (CB) and mature fruit (MF).

Rainfall, fertilizer applications, temperature, and irrigation hours fluctuated across our sampling period (Supplementary Fig. S1, Supplementary Table S3). Rainfall was sparse in this sample location (the Central Valley of California), ranging from 0.00 – 2.55 inches each month (Supplementary Fig. S1b, Supplementary Table S3) and total rainfall was a minor determinant of community structure across all four communities, explaining only 0.9 - 3.2% of the variation (PERMANOVA, *P* ≤0.001 - 0.104, *r*^*2*^ = 0.009 - 0.032) (Table 1). Similarly, fertilizer application describes a small percentage (1.0 - 5.8%) of the variation in the data for all four communities examined. We evaluated temperature based on the average temperature, and interactive effects it might have with water availability (Hrs. of irrigation and total rain) in order to capture the full range of conditions that could impact microbial community composition. Temperature had a minor impact on communities, as this factor only describes 0.5 - 4.2% of the variation in the data that include temperature as an interaction factor. In addition to phenology, interactions between phenology and sample year were a driving factor of leaf bacterial community composition, explaining 10.6% of the changes across the data (PERMANOVA, *P* ≤0.001, *r*^*2*^ = 0.106).

**Table 1.**
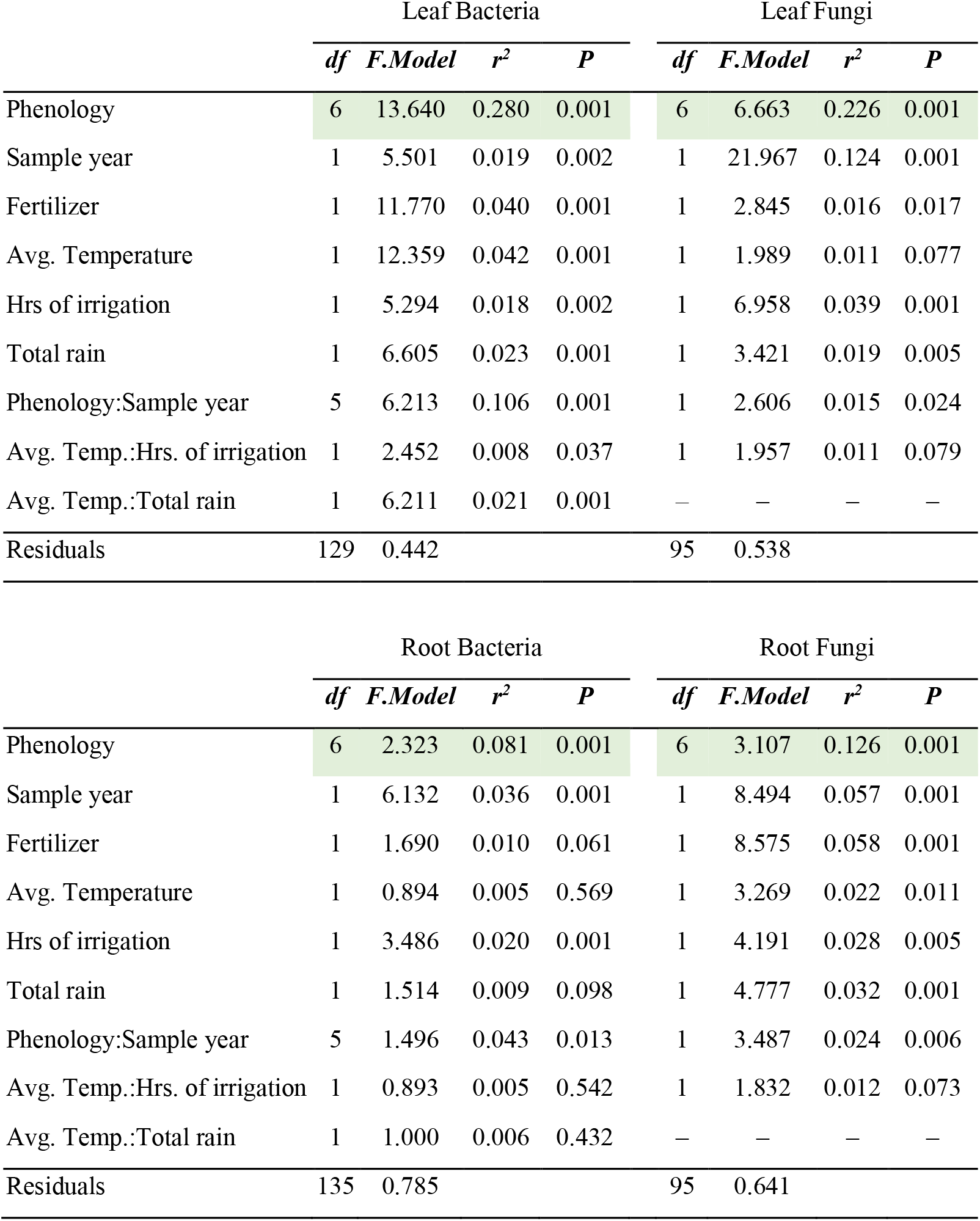
Beta diversity analysis (PERMANOVA) results for all variables. Test was performed using the vegan::adonis function and weighted UniFrac distances. Green highlights indicate the variable with the greatest r^2^ value (phenology).

|  | Leaf Bacteria |  |  |  | Leaf Fungi |  |  |  |
| --- | --- | --- | --- | --- | --- | --- | --- | --- |
| | *df* | *F.Model* | $r^2$ | *P* | *df* | *F.Model* | $r^2$ | *P* |
| Phenology | 6 | 13.640 | 0.280 | 0.001 | 6 | 6.663 | 0.226 | 0.001 |
| Sample year | 1 | 5.501 | 0.019 | 0.002 | 1 | 21.967 | 0.124 | 0.001 |
| Fertilizer | 1 | 11.770 | 0.040 | 0.001 | 1 | 2.845 | 0.016 | 0.017 |
| Avg. Temperature | 1 | 12.359 | 0.042 | 0.001 | 1 | 1.989 | 0.011 | 0.077 |
| Hrs of irrigation | 1 | 5.294 | 0.018 | 0.002 | 1 | 6.958 | 0.039 | 0.001 |
| Total rain | 1 | 6.605 | 0.023 | 0.001 | 1 | 3.421 | 0.019 | 0.005 |
| Phenology:Sample year | 5 | 6.213 | 0.106 | 0.001 | 1 | 2.606 | 0.015 | 0.024 |
| Avg. Temp.:Hrs. of irrigation | 1 | 2.452 | 0.008 | 0.037 | 1 | 1.957 | 0.011 | 0.079 |
| Avg. Temp.:Total rain | 1 | 6.211 | 0.021 | 0.001 | – | – | – | – |
| Residuals | 129 | 0.442 |  |  | 95 | 0.538 |  |  |

|  | Root Bacteria |  |  |  | Root Fungi |  |  |  |
| --- | --- | --- | --- | --- | --- | --- | --- | --- |
| | *df* | *F.Model* | $r^2$ | *P* | *df* | *F.Model* | $r^2$ | *P* |
| Phenology | 6 | 2.323 | 0.081 | 0.001 | 6 | 3.107 | 0.126 | 0.001 |
| Sample year | 1 | 6.132 | 0.036 | 0.001 | 1 | 8.494 | 0.057 | 0.001 |
| Fertilizer | 1 | 1.690 | 0.010 | 0.061 | 1 | 8.575 | 0.058 | 0.001 |
| Avg. Temperature | 1 | 0.894 | 0.005 | 0.569 | 1 | 3.269 | 0.022 | 0.011 |
| Hrs of irrigation | 1 | 3.486 | 0.020 | 0.001 | 1 | 4.191 | 0.028 | 0.005 |
| Total rain | 1 | 1.514 | 0.009 | 0.098 | 1 | 4.777 | 0.032 | 0.001 |
| Phenology:Sample year | 5 | 1.496 | 0.043 | 0.013 | 1 | 3.487 | 0.024 | 0.006 |
| Avg. Temp.:Hrs. of irrigation | 1 | 0.893 | 0.005 | 0.542 | 1 | 1.832 | 0.012 | 0.073 |
| Avg. Temp.:Total rain | 1 | 1.000 | 0.006 | 0.432 | – | – | – | – |
| Residuals | 135 | 0.785 |  |  | 95 | 0.641 |  |  |

Taken together, these beta diversity analyses indicate that plant phenological stage was the major driving factor in community composition for bacterial and fungal communities associated with leaves and roots. Significant compositional shifts are also visible at the phyla-level, particularly in the leaf bacterial community (Supplementary Fig. S3). Other covariates tested (irrigation, mean temperature, fertilizer applications, rainfall, and sample year) were minor or insignificant contributors to citrus-associated leaf and root microbiome composition.

### Stable, phylogenetically conserved microbial signatures across phenophases

We identified core microbial taxa for each of our seven phenological stages. Our core bacterial and fungal leaf and fungal root microbiomes include genera that were greater than 0.01%, and core root bacteriome greater than 0.1%, relative abundance in at least 75% of the samples within a phenological stage. All of our downstream analyses use genera that meet our core taxa cutoffs in at least one phenophase. We assessed our core taxa and separated them into three categories: (1) High stability=core member of six or more phenophases; (2) Medium stability=core member of three, four, or five phenophases; (3) Low stability=core member of two or less phenophases. We determined that of the identified core there were 3 (5.2%) leaf bacterial, 8 (30.7%) leaf fungal, 62 (70.4%) root bacterial, and 22 (61.1%) root fungal core genera that had high stability across phenophases (Fig. **4**, Supplementary Fig. S4). This suggests that both bacterial and fungal root communities have a substantially greater number of consistent or stable microbial features across the developmental cycle. However, our experimental design did not differentiate between endophytes versus epiphytes and, thus, may have missed some fine resolution microbial community shifts occurring between the endosphere and episphere. There were two bacterial (*Pseudomonas* and *Sphingomonas*) and one fungal (*Aureobasidium*) genera that were highly stable in both roots and leaves (Fig. **4c,d**).

**Figure 4.**
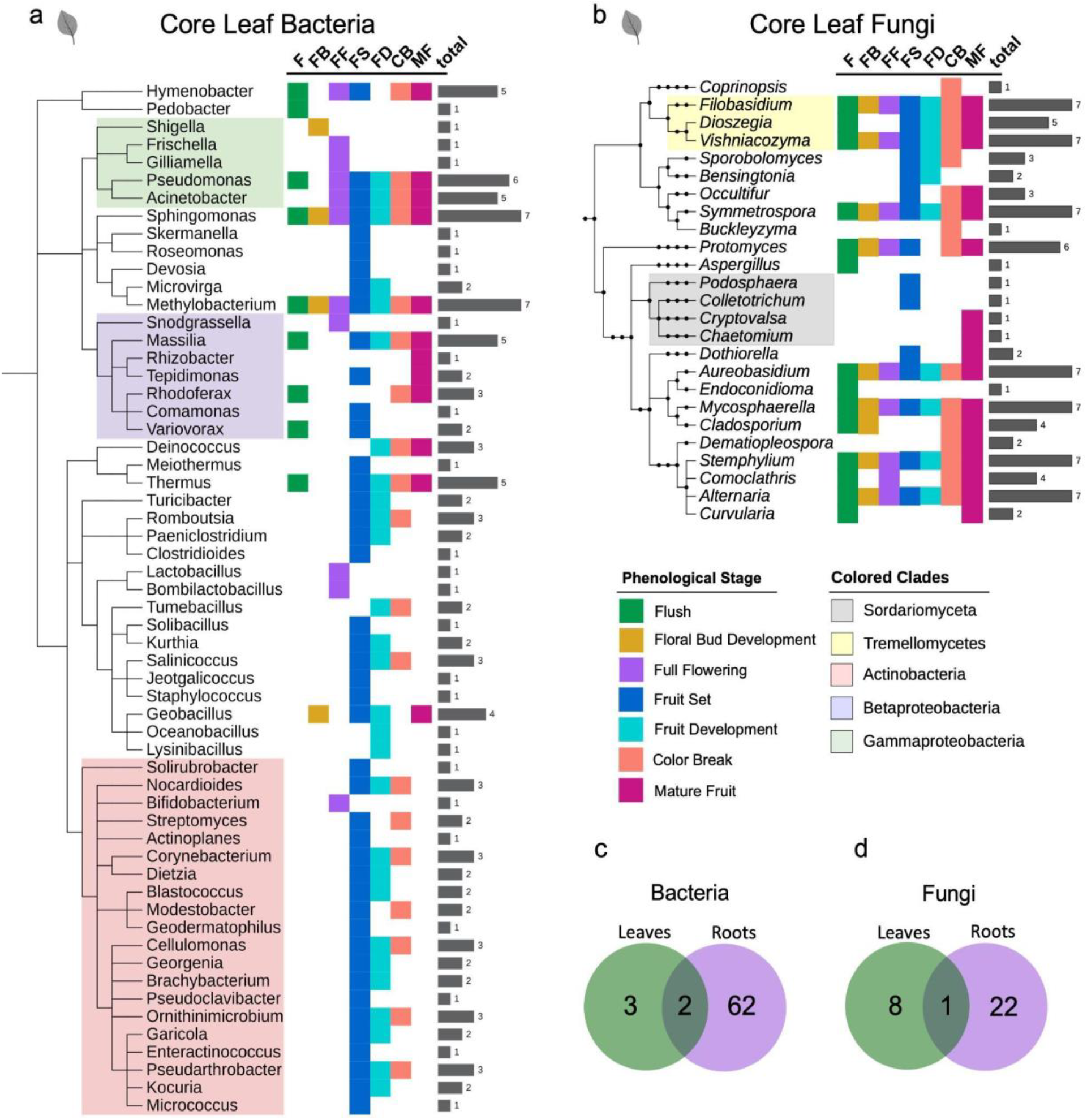
A subset of leaf bacterial and fungal core genera have phylogenetically conserved phenological association patterns. Phylogenetic trees of leaf bacterial (a) and fungal (b) genera that are core to one or more stages. Colored squares indicating a genus is core to flush (green), floral bud break (gold), full flowering (purple), fruit set (blue), fruit development (light blue), color break (salmon), and/or mature fruit (magenta). Gray bars indicate the total number of phenological stages each genus is core during. Venn diagrams show the number of highly stable core bacterial (c), and fungal (d) genera in leaf and root communities.

A phylogenetic analysis of the core genera indicates that both bacterial and fungal root communities were rich in highly stable and phylogenetically diverse core taxa (Supplementary Fig. S4). Root core genera from the bacterial clade Alphaproteobacteria (Class) and the fungal Family Pleosporomycetidae were all or nearly all binned as highly stable, indicating that genera in these clades were consistently high in relative abundance across all phenological stages. Medium and low stability core genera appear randomly dispersed across the root community phylogeny, with no obvious patterns.

However, leaf bacterial and fungal core community phylogenetic trees contained high, medium, and low stability patterns at the Class and Phyla levels (Fig. **4a,b**). All core genera in the fungal Class Tremellomycetes had medium to high stability. In contrast, all core genera in the fungal Class Sordariomycetes had low stability across phenophases, and only met the defined core cutoffs during fruit set or mature fruit stages. The leaf taxa within the bacterial Class Gammaproteobacteria consisted of genera with high, medium, and low stability across the phenophases. Interestingly, all the Gammaproteobacteria were a core member of the full flowering or floral bud break microbiomes regardless of their stability in other phenophases.

Another distinct phylogenetic pattern observed in the leaf community was genera in the bacterial Phylum Actinobacteria that had low or medium stability across all phenophases. However, 95.0% of core genera in the Actinobacteria clade were core during fruit set and/or fruit development. The only exception to this within the Actinobacteria clade was *Bifidobacterium*, which was associated only with full flowering and was not a core member of fruit set or fruit development microbiomes (Fig. **4a**). Lastly, the leaf bacterial Class Betaproteobacteria contains low to medium stability core genera with the most dispersed stage associations.

Overall, these data indicate that root bacterial and fungal communities have greater stability across phenophases than leaves (Fig. **4c,d**). Additionally, core taxa had phylogenetically related trends within the high, medium, and low stability classifications indicating that conserved, vertically descended microbial traits may play a role in determining bacterial and fungal associations across phenophases, particularly in above ground leaf tissue.

### Specific taxa were enriched in the foliar microbiome across the flowering phenophases

We completed a genus-level differential abundance analysis on our list of core taxa that were ≥0.01% relative abundance and ≥75% prevalence in one or more phenophases. Our differential abundance analysis can determine finer scale phenophase associations beyond just classification as a core microbiome member by looking for increases in abundance (enrichments) during specific phenophases. Among all the phenophases, those associated with flowering (floral bud development and full flowering), had striking microbial enrichments, particularly among the leaf bacteria. *Acinetobacter* was a core member of five phenophases, but was significantly enriched during full flowering when compared to other phenophases (Fig. **5a**). *Acinetobacter* had a gradual enrichment from flush, floral bud development, to full flowering. This gradual enrichment signature indicates that *Acinetobacter* was present throughout the year, but has a high temporal turnover rate that is in sync with the transitions from flush to floral bud development and then to full flowering.

**Figure 5.**
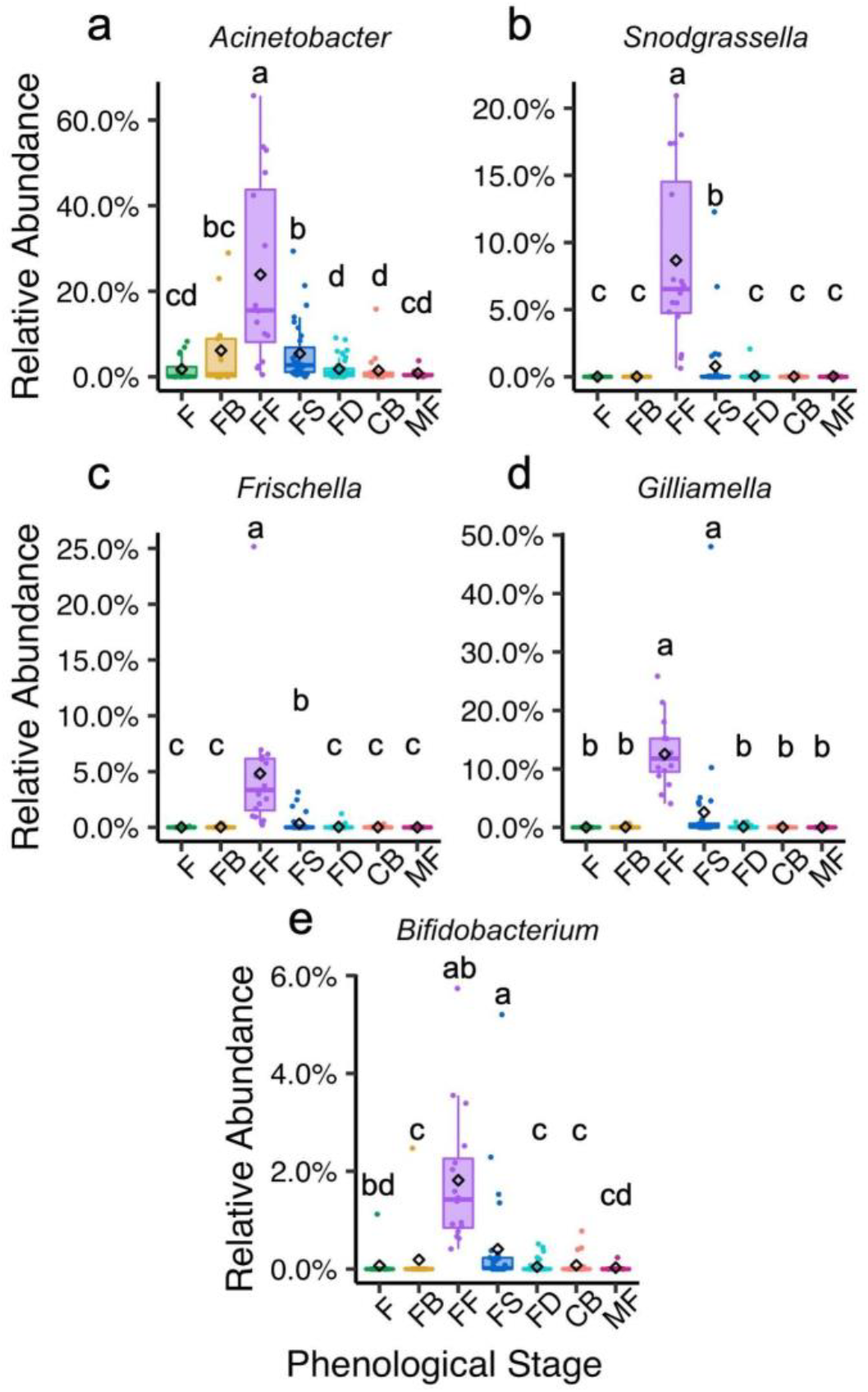
Core leaf bacteriome members enriched during full flowering. Gradually (a-b) and suddenly (c-e) enriched taxa during full flowering. The diamond symbol indicates the mean relative abundance. Letters indicate significant differences of *P* ≤ 0.05, determined using DESeq2 GLM, Wald test with FDR adjustment. Phenological stages on the x-axis include: flush (F), floral bud break (FB), full flowering (FF), fruit set (FS), fruit development (FD), color break (CB) and mature fruit (MF).

We also observed bacteria that were sharply enriched during full flowering rather than undergoing gradual enrichments over the phenophases that lead up to full flowering (flush and floral bud development). These include *Snodgrassella, Frischella, Gilliamella*, and *Bifidobacterium* (Fig. **5b-e**). The sharp enrichment patterns during full flowering suggest that these taxa are introduced into the community via a dispersal event.

### Foliar microbial depletions associated with flowering

We also identified bacterial leaf genera that had significant depletions during floral bud development and/or full flowering (Fig **6a-h**). Four Actinobacter genera *Corynebacterium, Dietzia, Georgenia*, and *Ornithinimicrobium* were significantly depleted during floral bud development and full flowering (Fig. **6a-d**). *Bacillus, Methylobacterium, Romboutsia*, and *Sphingomonas* also significantly decreased in relative abundance during floral bud development and/or full flowering (Fig. **6e-h**). For all differentially abundant genera, including bacteria and fungi, across all phenophases see supplemental figure S5, and supplemental table S4.

**Figure 6.**
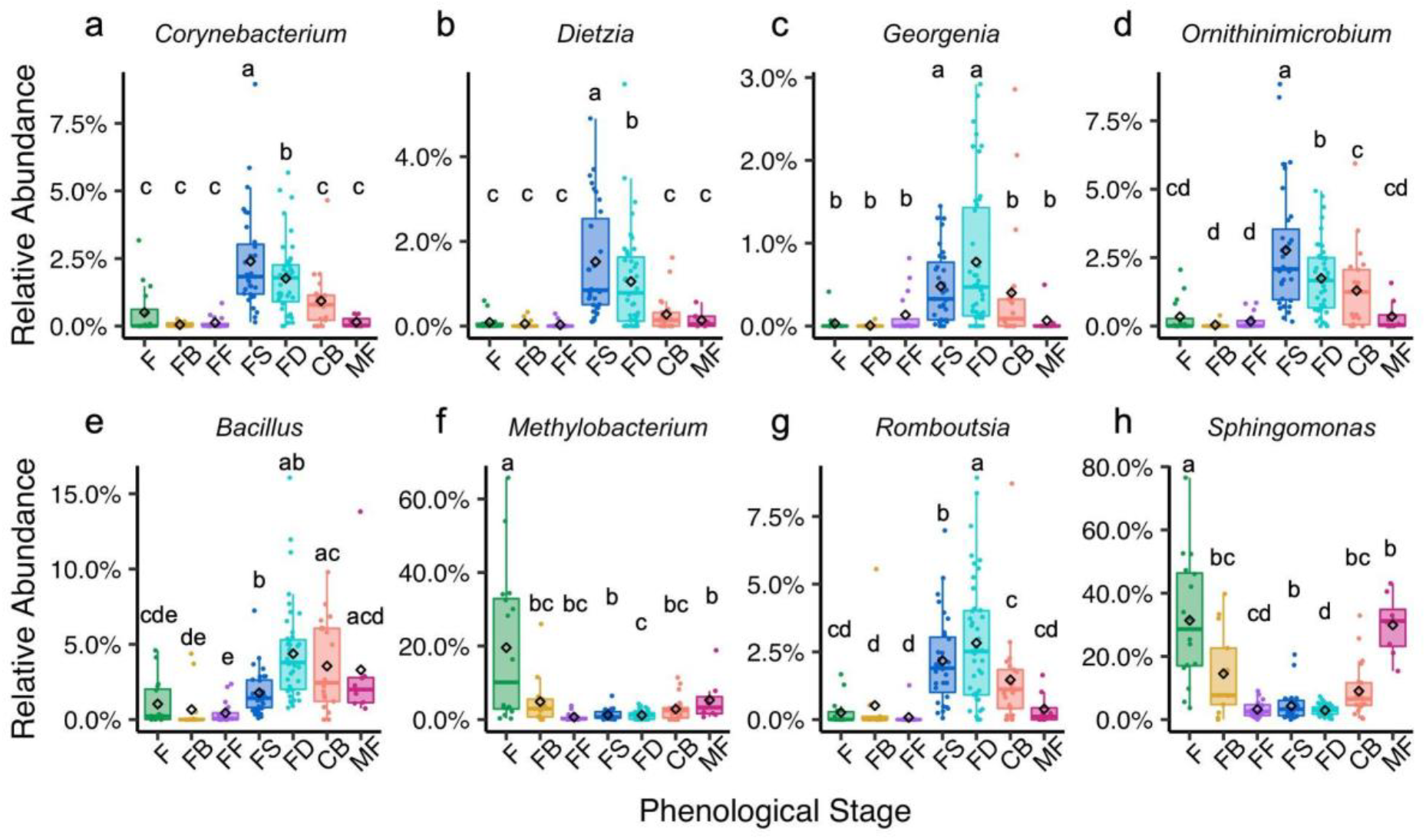
Core leaf bacteriome members depleted during full flowering. The diamond symbol indicates the mean relative abundance. Letters indicate significant differences of *P* ≤ 0.05, determined using DESeq2 GLM, Wald test with FDR adjustment. Phenological stages on the x-axis include: flush (F), floral bud break (FB), full flowering (FF), fruit set (FS), fruit development (FD), color break (CB) and mature fruit (MF).

### Microbe-microbe interactions contribute to phenophase specific community structure

We performed a network analysis on the foliar bacterial communities from all samples with a focus on the significantly enriched and/or depleted populations and any populations they have direct connections with (neighbors). The goal of this approach was to give a broad overview of bacterial interactions across phenophases and identify taxa that potentially interact with specific phenophase-enriched taxa and potentially play a role in observed seasonal community compositional shifts. *Rhizobium, Sphingomonas*, an unknown bacteria, an unknown Bacillaceae (family), *Acinetobacter*, and *Romboutsia* have the highest normalized betweenness centrality scores ranging from 0.110 – 0.187. Betweenness centrality is a proxy for influence within a network because it measures how often a particular node (i.e., taxon) is the shortest connection or bridge between two other nodes. These high betweenness centrality scores and placement within the network indicates that these genera are potentially keystone taxa that may perform a stabilizing role in the microbial communities across phenological transitions and events (Fig. **7**, red nodes). Groups of taxa connected by putative positive interactions cluster together to form distinct modules. These modules are separated by putative negative interactions. Our analysis organized bacterial taxa that were enriched in fruit set and fruit development into a single highly connective community module (cluster) (Fig. **7**, blue nodes). This suggests that fruit set and fruit development associated microbiomes are compositionally similar and few microbe-microbe interactions change during the transition from fruit set to fruit development. Leaf bacteria associated with flowering also formed a module within the network (Fig. **7**, purple nodes). Specific bacteria within the fruit set/development and flowering modules also interact with taxa that were enriched in the other four phenophases, which cluster together into a third module (Fig. **7**, grey nodes). Overall, these predicted positive interactions represent inter- or co-dependent microbe-microbe relationships, and the putative negative interactions indicate potential direct (e.g., antibiosis) or indirect (e.g., resource exclusion) competition. These predicted microbe-microbe interactions within the microbiome likely impact community composition in addition to the exogenous influences of abiotic environmental conditions and biotic host physiological factors (e.g., carbon availability).

**Figure 7.**
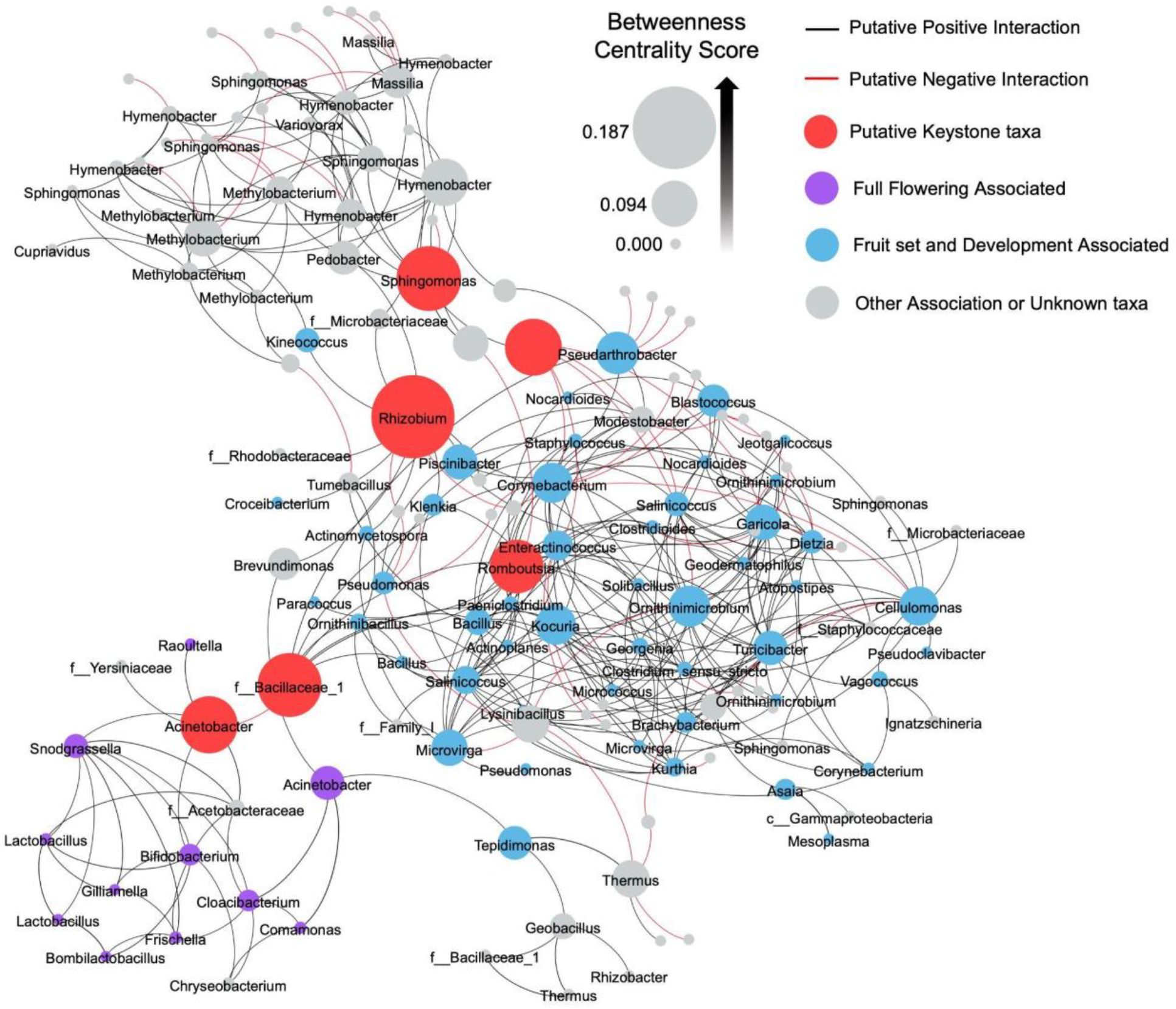
Core leaf bacteriome interaction network. Each node represents a leaf bacteria OTU and is labeled with the lowest known taxonomic rank. Nodes are sized by betweenness centrality scores calculated using Gephi. Red nodes are predicted keystone taxa. Purple nodes are taxa significantly enriched during full flowering (FF), blue colored nodes are taxa significantly enriched during fruit set (FS) and/or fruit development (FD), and grey colored nodes are not significantly enriched during FF, FS, or FD. Lines represent predicted positive (black) and negative (red) interactions, determined using SPIEC-EASI sparse neighborhood covariance selection to infer interactions.

## DISCUSSION

The majority of studies examining how plant developmental stage impacts the plant’s microbiome have focused on bacteria associated with the rhizosphere of annuals or herbaceous perennials such as maize^43,44^, rice^28^, sorghum^26,45^, wheat^46^, *Arabidopsis*^29^ and *Boechera*^30^. These important studies indicate that rhizosphere-associated microbiomes can shift in association with plant developmental stages in both domesticated and wild plants that have short-lived above ground tissues. Studies of the endophytic xylem sap microbiome in grapevine, a deciduous perennial, also showed microbial shifts were linked to changes in phenological stage^47^. However, much less is known about how overall plant phenology impacts above and below ground microbiomes of evergreen woody perennials that have lifespans that can be decades long and can retain their leaves for multiple years, as compared to annuals or deciduous perennials that produce and shed all their leaves each season. Here, we investigated microbiome dynamics in above and below ground tissues of mature twenty-year old *Citrus sinensis* trees to determine if temporal microbiome fluctuations were associated with host phenological events. The unique contribution of our research was the separation of leaf development from tree phenology. We did this by analyzing the changes in the foliar microbiome on fully mature leaves, which developed as part of the same leaf cohort from the previous year, in relation to the phenological stages of the current year. Thus, are exposed to the same starting inoculum, minimizing the bias of any potential priority effects (i.e., order of arrival).

Our results indicate that the phyllosphere microbiome has an active and dynamic relationship with host phenology. More specifically, microbial shifts occurred as trees transitioned from the spring leaf flushing stage and entered flowering. The transition from spring flush to floral bud development and full flowering aligns with important transitions in source-to-sink transport of photosynthate in the tree^4^. During foliar flushing periods, young leaves are a primary carbohydrate sink as they rapidly expand and mature. This source-to-sink transport of photosynthate essentially reverses during floral bud break and development, when mature leaves transition to serve as source tissues and begin transporting photosynthates to developing floral tissues that are now the primary sink tissues. In addition to changes in source-to-sink transport, there are also significant changes in water dynamics within the canopy of the tree associated with full flowering. Flowers have the highest transpiration rate of the tree even compared to the leaves, which drastically increases the amount of water being transported into the overall canopy of the tree^4^. Interestingly, the significant shift in overall foliar community composition from flushing to full flowering was not coupled with a change in species richness indicating that the same taxa were present, just in different relative abundances in relation to one another. This demonstrates that foliar microbiome assemblage is changing in sync with tree physiology and development.

Empirical data, including presence/absence and relative abundance, can also be used to infer patterns or microbial enrichments and/or depletions as well as ecological mechanisms that contribute to plant microbiome assembly, such as microbial species turnover and dispersal^48–50^. Interestingly, microbial enrichment and depletion patterns of specific taxa suggest that microbial species turnover and dispersal events within the citrus microbiome occur in sync with phenological stage transitions. These enrichment/depletion patterns for specific taxa were more apparent in leaves than in the root compartment. Specifically, the bacterial genus *Acinetobacter* was enriched in leaves as trees transitioned from spring flush to floral bud development and peaked in relative abundance during full flowering, which is when leaves shift from acting as sink tissues to becoming source tissues. This may create a microenvironment that selects for an increase in relative abundance of these taxa when carbohydrate is translocating out of the leaves. Plant-associated *Acinetobacter* spp. have plant growth promoting properties that include antagonism towards fungi^51^, the ability to solubilize phosphate and to produce the plant hormone, gibberellic acid^52,53^. *Acinetobacter* spp. are highly abundant in the floral nectar microbiome of *Citrus paradisi* and other plant species^54,55^ and were identified as a core member of the grapevine xylem sap microbiome^47^. Its significant increase in relative abundance in the leaf microbiome at the time of flowering in citrus suggests a potential synergy between the foliar and floral microbiomes. *Acinetobacter* was also predicted to be a keystone taxon and was a major link between the flowering community and fruit set/development community clusters in our network. This enrichment in *Acinetobacter* may be simply due to selection imposed on the microbial community by the local plant environment, but it is tempting to speculate that *Acinetobacter* spp. provide an exogenous service to the plant by producing gibberellic acid and biologically available phosphorus to promote flowering that is in phase with its host’s phenological development. This hypothesis that the plant environment selects for taxa within its foliar microbiome that, in turn, promotes its own reproductive growth warrants future inquiry. Specific bacterial enrichments also occurred at bloom time in grapevine further supporting the evidence of a host driven microbial response to environmental cues derived from shifts in plant developmental stage^47^.

We also observed signatures that indicate specific taxa were depleted in relative abundance during flowering, but enriched during fruit set. Phylogenetic reconstruction of these taxa indicates that the majority of the taxa belonging to the Actinobacteria phylum (19 of the 20 genera) were significantly depleted during flowering, but subsequently enriched when trees begin to set fruit. This phylogenetic conservation of depletion/enrichment patterns within the Actinobacteria clade indicates that this is a non-random fluctuation within the microbiome structure associated with the transition from flowering to fruit production. As citrus trees set fruit, the fruits themselves begin exporting and importing hormones, such as indole acetic acid and cytokinins, respectively^3^. This results in a change in hormone levels in leaves as well. These hormonal shifts may place selective pressure on the microbial community that the foliar Actinobacteria are particularly responsive to that lead to significant enrichments during fruit set and development. Specific differentially abundant taxa within the Actinobacteria clade that followed this pattern included *Corynebacterium, Dietzia, Georgenia*, and *Ornithinimicrobium*. Members of these genera can fix nitrogen and produce IAA, both of which are important supporters of fruit development^56–59^. Thus, it is tempting to speculate that these taxa could play a role in co-regulating fruit development in a manner that is synergistic with the host’s production of reproductive hormones. The biological role of Actinobacteria in the foliar microbiomes of plants is not well understood but overall species richness was conserved across all phenological stages, except for fruit set, indicating that this phenological stage allows for microbial enrichments of specific taxa, particularly those belonging to the Actinobacteria phylum. Leaf bacteria within the same clade may have similar functional roles, suggesting that members of the *Actinobacteria* phylum play an important role in the foliar microbiome during fruit set.

Genera outside of the Actinobacteria clade were also depleted in leaves during floral bud development and full flowering, including *Bacillus, Methylobacterium, Romboutsia*, and *Sphingomonas*. Notably, *Romboutsia* and *Sphingomonas* were predicted to be keystone taxa in our microbe-microbe interaction network analysis, and all are in the top 20% highest betweenness centrality scores. Keystone taxa play a stabilizing role in microbial communities. Depletion of these taxa during flowering may have cascading effects that influence microbial species turnover by allowing other taxa, such as the Actinobacteria, to flourish during subsequent developmental stages like fruit set. This suggests that microbial turnover in the foliar microbiome is mediated by selective pressures imposed by the plant developmental stage in conjunction with microbe-microbe interactions to modulate community diversity and composition.

Microbial dispersal events can drive microbial turnover and influence the relative abundance of endogenous taxa in the community. Full flowering is a dynamic phenophase in plant development where there are frequent interactions between plants and pollinator species that rely on floral resources, like nectar and pollen. These macro-level interactions can also have impacts at the microorganismal level. Pollinator (e.g., bee) visitation alters flower surface, nectar, and subsequent seed microbial community composition^60–63^. During flowering, we observed sudden microbial enrichments of bacteria taxa belonging to the Betaproteobacteria and Gammaproteobacteria clades that include *Giliamella, Snodgrassella, Bifidobacterium*, and *Frischella*. These enrichments in these anaerobic taxa were unique to the flowering phenophase and quickly declined following flowering, suggesting they are immigrants to the community and not endogenous members of the native microbiome. Moreover, these anaerobic taxa are prevalent in the bee gut microbiome^64^, therefore we hypothesize that these taxa were dispersed into the citrus microbiome during honeybee visitation. We consider this external influence host phenology associated because phenophase specific plant morphology (i.e., flowers) regulate this diffuse interaction. Bacteria can be introduced to plants by bees and potentially migrate from the flower to the vascular bundles resulting in systemic movement within the plant^65–67^. Leaf carbohydrate content is the highest during flowering, which may promote the growth of these fermenting bacteria^4^. Notably, *Bifidobacterium* was the only core leaf genus from the Actinobacteria phylum that was enriched during flowering whereas the other 19 Actinobacteria taxa were depleted during flowering. This further supports the hypothesis that *Bifidobacterium* was introduced via a dispersal event and is not part of the endogenous microbiota like the other taxa in the plant-associated Actinobacteria clade. Nectar-inhabiting bacteria can influence nectar volatile profiles that, in turn, influence pollinator visitation preferences^68^ and it would be interesting to determine if these putative immigrants contribute to shifts in nectar volatile profiles that affect bee feeding behaviors.

The next frontier in microbiome research is to determine the functional roles that microbes play in microbe-microbe and host-microbiome interactions. Martiny et al. 2013 found that conservation of microbial traits was more strongly linked to vertical phylogenetic relatedness of the microorganisms within a microbiome than to traits that are shared among taxa by horizontal gene transfer^69^. Similarly, we also observed phylogenetic conservation within microbial enrichments suggesting that those groups of related organisms play similar functional roles during specific phenophases or across several phenophases. We speculate that taxa with high stability across phenophases may serve a community stabilizing function, while low stability or phenophase specific core microbes likely have more specialized, transient roles in the community. Because microbes can alter host phenology^17,20,70–73^, which is a critical factor in plant health and productivity, incorporation of microbial presence/absence and patterns of enrichment into plant phenological models may improve phenophase timing prediction, once the functional roles of these microbes are determined. This information could also lead to the commercialization of biofertilizers for horticultural purposes that could be applied at specific plant life stages to enhance crop productivity.

## ACKNOWLEDGMENTS

The authors would like to thank Drs. Carol Lovatt, Kurt Anderson, and Nicole Rafferty for their insightful comments. This work was supported by USDA NIFA Grant No. 2017-70016-26053, CDFA Grant No. SCB16056 and 19-0001-034-SF, USDA National Institute of Food and Agriculture Hatch Projects 1002710, 233883 and the National Science Foundation Graduate Research Fellowship Program under Grant No. (NSF DGE-1326120). Any opinions, findings, and conclusions or recommendations expressed in this material are those of the author(s) and do not necessarily reflect the views of the National Science Foundation.

## COMPETING INTERESTS

The authors declare that they have no competing interests.

## AUTHOR CONTRIBUTIONS

NAG and MCR designed the project. NAG and PR conducted fieldwork. NAG and IDA processed field samples. NAG and FCFV built sequencing libraries. NAG performed data analyses. NAG and MCR wrote the manuscript. All authors contributed to the editing of the manuscript.

## DATA AVAILABILITY

The data that supports the findings of this study are openly available in the NCBI SRA database under BioProject PRJNA685913. Data is available for review at the following link: https://dataview.ncbi.nlm.nih.gov/object/PRJNA685913?reviewer=6s46aglvstjai5arhqs2cp7434

## SUPPLEMENTAL MATERIAL LEGENDS

**Supplementary Table S1. Bacterial 16S read counts**.

**Supplementary Table S2. Phenological stage Pairwise-PERMANOVA results (adjusted *P* values)**. Significant differences in beta diversity (*P* ≤ 0.05) are indicated by bolded font. Phenological stages include: flush (F), floral bud break (FB), full flowering (FF), fruit set (FS), fruit development (FD), color break (CB) and mature fruit (MF).

**Supplementary Table S3. Metadata file**.

**Supplementary Table S4. All significant differentially abundant bacterial and fungal genera across phenological stages and tissue types**.

**Supplementary Figure S1. Environmental factors fluctuate during citrus development**. The primary phenophase displayed during each month from July 2017 to April 2019 is indicated by colors on the x-axis. a) gray bars represent total hours of irrigation each month. b) Navy bars represent mean total rainfall each month. c) Points indicate average high (red), average low (blue), and total monthly average (black) temperatures.

**Supplementary Figure S2. Sample year has minor effects on community diversity and composition**. Beta diversity PCoA plots of bacterial leaf (a) and bacterial root (b) communities. Points are colored by sample year and represent a complete community from a single leaf or root sample. Ellipses represent 95% confidence intervals.

**Supplementary Figure S3. Phyla level compositional changes across phenological stages**. Stacked bar plots showing relative abundance of bacterial leaf (a), fungal leaf (b), bacterial root (c), and fungal root (d) phyla across phenological stages. Phenological stages on the x-axis include: flush (F), floral bud break (FB), full flowering (FF), fruit set (FS), fruit development (FD), color break (CB) and mature fruit (MF).

**Supplementary Figure S4. Phenological Core root bacterial and fungal genera**. Phylogenetic trees of root bacterial (a) and fungal (b) genera that are core to one or more stages. Colored squares indicating a genus is core to flush (green), floral bud break (gold), full flowering (purple), fruit set (blue), fruit development (light blue), color break (salmon), and/or mature fruit (magenta). Gray bars indicate the total number of phenological stages where that genus is core.

**Supplementary Figure S5. Core leaf mycobiome members with significant enrichments**. The diamond symbol represents the mean relative abundance. Letters indicate a significant difference of *P* ≤ 0.05, determined using DESeq2 GLM, Wald test with FDR adjustment. Phenological stages on the x-axis include: flush (F), floral bud break (FB), full flowering (FF), fruit set (FS), fruit development (FD), color break (CB) and mature fruit (MF).

